# SIR proteins create compact heterochromatin fibers

**DOI:** 10.1101/346296

**Authors:** Sarah G. Swygert, Subhadip Senapati, Mehmet F. Bolukbasi, Scot A. Wolfe, Stuart Lindsay, Craig L. Peterson

## Abstract

Heterochromatin is a silenced chromatin region essential for maintaining genomic stability and driving developmental processes. The complicated structure and dynamics of heterochromatin have rendered it difficult to characterize. In budding yeast, heterochromatin assembly requires the SIR proteins -- Sir3, believed to be the primary structural component of SIR heterochromatin, and the Sir2/4 complex, responsible for the targeted recruitment of SIR proteins and the deacetylation of lysine 16 of histone H4. Previously, we found that Sir3 binds but does not compact nucleosomal arrays. Here we reconstitute chromatin fibers with the complete complement of SIR proteins and use sedimentation velocity, molecular modeling, and atomic force microscopy to characterize the stoichiometry and conformation of SIR chromatin fibers. In contrast to previous studies, our results demonstrate that SIR arrays are highly compact. Strikingly, the condensed structure of SIR heterochromatin fibers requires both the integrity of H4K16 and an interaction between Sir3 and Sir4. We propose a model in which two molecules of Sir3 bridge and stabilize two adjacent nucleosomes, while a single Sir2/4 heterodimer binds the intervening linker DNA, driving fiber compaction.

## Introduction

Eukaryotic cells regulate the accessibility of their genome to enzymatic processes by organizing it into two functionally distinct compartments, known as euchromatin and heterochromatin. Euchromatin consists of actively transcribed gene regions, whereas heterochromatin is refractory to external processes such as transcription and recombination ^1^. Heterochromatin organizes and protects centromeres and telomeres, guards against the spreading of transposons, and prevents aberrant homologous recombination within repetitive regions that can lead to chromosomal abnormalities such as deletions, inversions, and translocations.^2^^-^^4^ Additionally, heterochromatin formation is an essential developmental process that drives the differentiation and maintenance of cell types.^2,3,5^ Although heterochromatin carries a distinct subset of histone modifications and protein complexes, the mechanism by which heterochromatin maintains its silent state is poorly understood.

The most thoroughly characterized heterochromatin state exists in the budding yeast *Saccharomyces cerevisiae*, which requires the SIR proteins -- Sir2, Sir3, and Sir4 -- for silencing.^6,7^ The formation of SIR heterochromatin is believed to be a step-wise process in which a Sir2/4 complex is initially recruited to silencing regions via interactions between the Sir4 protein and sequence-specific DNA binding factors such as Rap1, Orc1, and Abf1.^8^^-^^13^ Sir2, an NAD^+^-dependent histone deacetylase, then deacetylates the H4 tail of an adjacent nucleosome at lysine 16 (H4K16), which promotes binding of the Sir3 protein to the nucleosome, in turn recruiting additional Sir2/4 complex.^10,14-16^ As this cycle of deacetylation and binding continues, the SIR proteins spread away from the nucleation site, creating a silent, heterochromatin domain.^17^

The importance of H4K16 to SIR heterochromatin was initially discovered when its mutation to glutamine (H4K16Q) was found to disrupt the repression of the silent mating loci, and compensatory mutations in the Sir3 protein were identified.^18^ The physical interaction between Sir3 and H4K16 has been explored at length both *in vivo* and *in vitro*,^14,19-21^ with several crystal structures of a Sir3-nucleosome complex displaying an electronegative patch of Sir3 that forms a binding pocket for H4K16.^22^^-^^24^ Interestingly, while Sir3 alone demonstrates a clear preference for binding unmodified versus acetylated or mutated H4K16, a purified Sir2/3/4 complex binds with nearly equal affinity to acetylated nucleosomes or nucleosomes harboring H4-K16Q.^25^^-^^27^ Notably, this impact of Sir2/4 on the binding specificity of Sir3 is lost when a single amino acid substitution is made within the Sir3 interaction domain of Sir4 (Sir4I1311N),^25,28,29^ suggesting that the binding of Sir3 to Sir4 modifies its affinity for nucleosomes that contain H4K16Ac or H4K16Q. Furthermore, although all three SIR proteins can bind to H4K16Ac chromatin templates, the resulting Sir chromatin fibers do not block transcription in the absence of NAD^+^.^25,26^ This suggests that although an interaction between modified H4K16 and SIR proteins is possible, the presence of H4K16Ac prevents the formation of a functional, repressive heterochromatin structure.

Previously, we established experimental conditions in which Sir3 displays a strong binding preference for reconstituted nucleosomal arrays that contained wildtype (WT) histones compared to arrays assembled with histones with the H4K16Q substitution.^21^ We found that these reconstituted Sir3 chromatin fibers remained quite extended compared to 30 nanometer (nm) fibers that were condensed with divalent cations. Here, we characterize heterochromatin fibers reconstituted with all three SIR proteins. Consistent with previous studies,^25^^-^^27^ we find that the full complement of SIR proteins binds with a similar stoichiometry to both WT and H4K16Q chromatin, but only with WT chromatin are SIR proteins competent to form a compacted structure that is consistent with an inaccessible heterochromatin fiber.

## Results

### Sir2/4 binds to both WT and H4K16Q arrays

Previously, we established optimal ionic conditions for the formation of Sir3 chromatin fibers that are sensitive to H4K16Q.^21^ In order to examine the binding of Sir2/4 to nucleosomal arrays under these same conditions, we assembled recombinant WT or H4K16Q nucleosomal arrays by salt dialysis using a DNA template containing twelve head-to-tail repeats of a 601 nucleosome positioning sequence (601-177-12). Binding of purified Sir2/4 complex was first analyzed by an electrophoretic mobility shift assay (EMSA), in which increasing amounts of Sir2/4 complex were added to nucleosomal arrays and binding was monitored by the decrease in mobility on an agarose gel (**Fig. 1a**). Consistent with similar studies,^27^ Sir2/4 bound to WT and H4K16Q arrays at similar concentrations, with an apparent slight preference for H4K16Q (**Fig. 1a**). Next, Sir2/4 was titrated onto these nucleosomal arrays and interactions were monitored by sedimentation velocity analysis in an analytical ultracentrifuge (SV-AUC). Sir2/4 bound maximally to arrays at a ratio of 1-2 Sir2/4 molecules per nucleosome (**Fig. 1a** and **b**). Beyond a molar ratio of 2 Sir2/4’s per nucleosome, arrays became insoluble and were not used in sedimentation experiments. Notably, Sir2/4 appears to interact equally well with both WT and H4K16Q arrays, sedimenting at 33-38S upon addition of 2 molecules of Sir2/4 per nucleosome. This contrasts with the sedimentation properties of Sir3 nucleosomal arrays that only assemble effectively with WT histones and sediment at 45-50 S (Swygert, 2014; see also **Fig. 2a** and Fig. **5a**).

**Figure 1.**
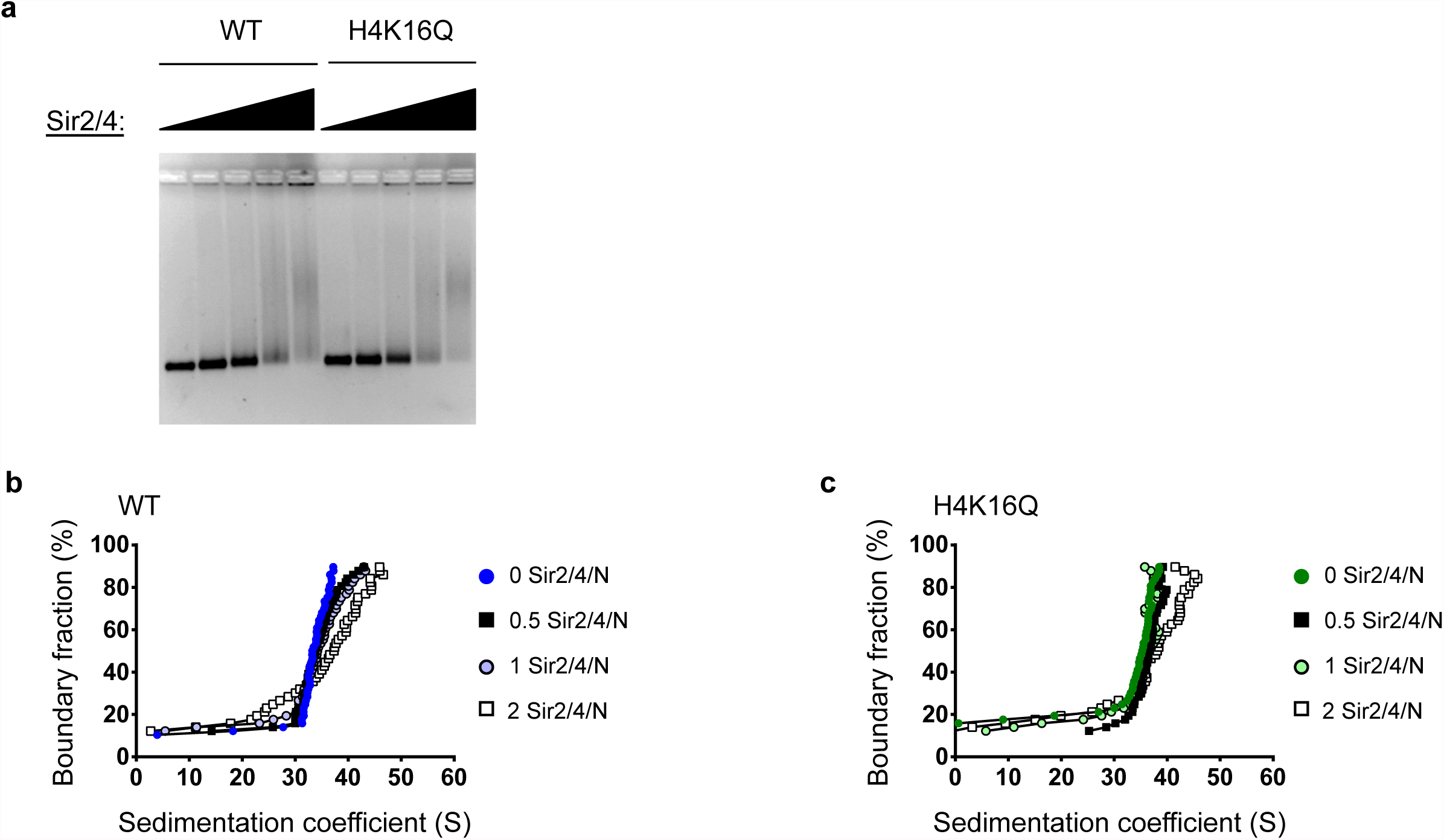
The Sir2/4 complex binds both WT and H4K16Q arrays. **(a)** EMSA of Sir2/4 binding to WT and H4K16Q arrays. **(b-c)** vHW plots of Sir2/4 complex titrated onto WT and H4K16Q arrays. Numbers indicate the molar ratio of Sir2/4 complex per nucleosome.

**Figure 2.**
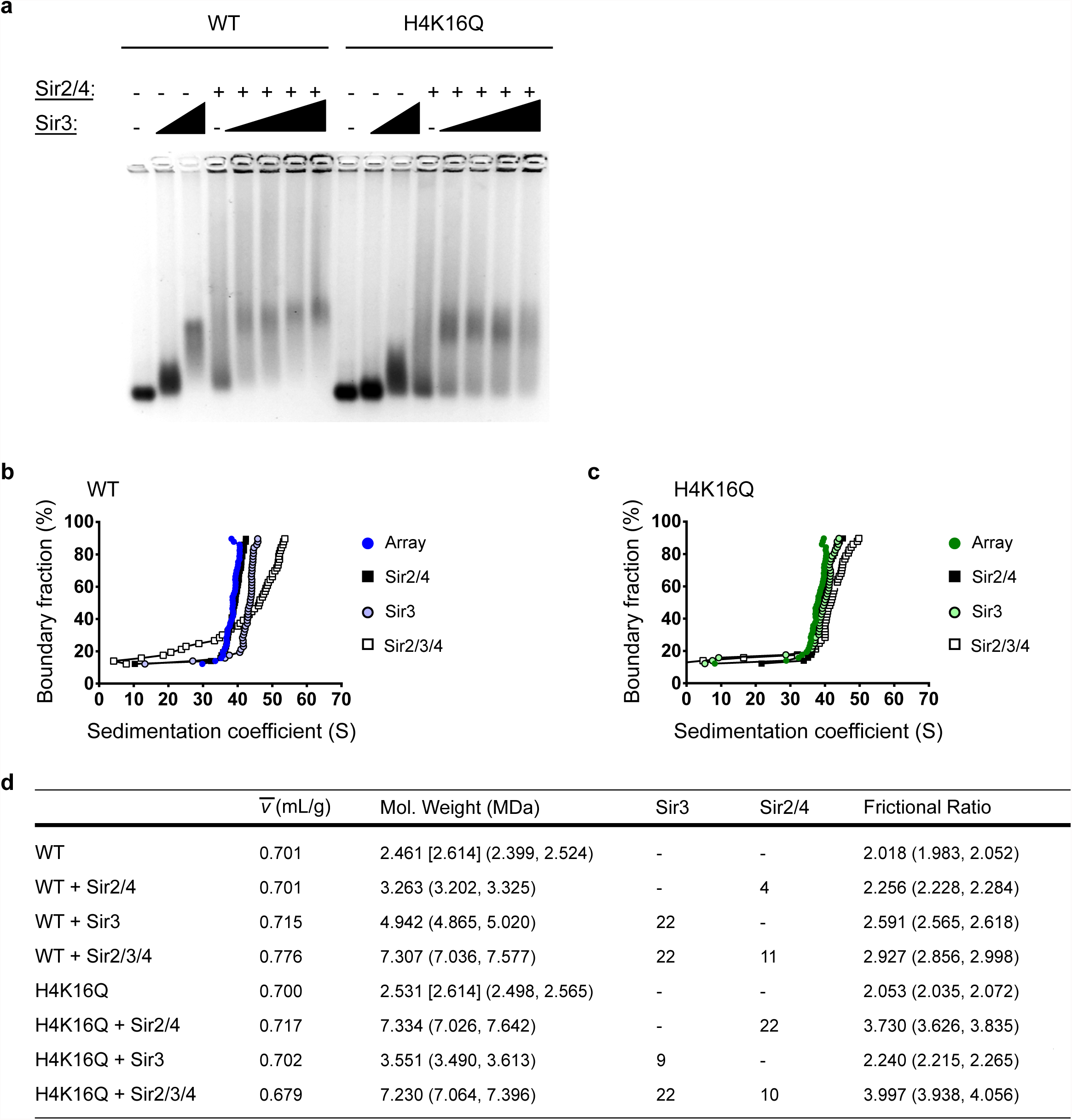
SIR interactions with WT and H4K16Q arrays are distinct. **(a)** EMSA of Sir3 titrated onto WT and H4K16Q arrays in the absence or presence of Sir2/4 complex. **(b-c)** vHW plots of Sir2/4, Sir3, and Sir3 and Sir2/4 complex added to WT and H4K16Q arrays. **(d)** 2DSA/GA-MC modeling results of the sedimentation data in **(b-c)**. Numbers in brackets represent expected molecular weights. Numbers in parentheses are 95% confidence intervals. Stoichiometries upon addition of Sir2/3/4 are speculative.

### SIR interactions with WT and H4K16Q arrays yield fibers with distinct solution dynamics

To investigate the binding properties of arrays reconstituted with all three SIR proteins, increasing amounts of Sir3 was added to reactions that contained nucleosomal arrays and Sir2/4, and binding was first evaluated by EMSA (**Fig. 2a**). In isolation, Sir3 bound with much greater affinity to WT than H4K16Q arrays, as seen previously. However, titrating Sir3 in the presence of Sir2/4 led to the formation of a complex with similar mobility for both WT and H4K16Q arrays. Notably, this complex remained at constant mobility despite further additions of Sir3, suggesting that the ionic conditions that have been used to form a discrete Sir3 fiber also promote formation of a discrete SIR heterochromatin fiber. This is in stark contrast to previous studies in different buffer conditions whereby addition of increasing concentrations of SIR proteins led to continual decreases in the mobility of EMSA species, suggestive of nonspecific DNA binding or aggregation.^27,30^

SIR heterochromatin fibers were next examined by SV-AUC (**Fig. 2b** and **c**). As in **Fig. 1**, the addition of Sir2/4 onto WT and H4K16Q arrays led to only small changes in the sedimentation coefficient (S) distribution. In contrast, addition of Sir3 to WT arrays led to a more substantial change in S (~35-45 S), and addition of Sir3 did not shift H4K16Q arrays at all, consistent with binding to wildtype but not H4K16Q nucleosomes. Strikingly, addition of all three SIR proteins to WT arrays led to a large increase in the sedimentation coefficient distribution (~50S), whereas binding of all three SIR proteins to H4K16Q arrays led to a modest shift to ~42 S. This result contrasts with the similar mobility of these fibers in the EMSA assay, and suggests that the binding of Sir proteins to WT and H4K16Q arrays may yield similar stoichiometries but distinct conformations in solution.

The sedimentation behavior of a macromolecule in an SV-AUC experiment is proportional to both its buoyant molecular weight and its frictional properties governed by its overall shape.^31^^-^^33^ Consequently, the observed SIR-induced changes in the S distribution of nucleosomal arrays in **Fig. 2** could be due to an increased molecular weight, an altered conformation of the nucleosomal fiber, or a combination of both. To separate these parameters, we applied a set of modeling methods, 2DSA/GA-MC, which fit the sedimentation coefficient, partial concentration, molecular weight, and frictional ratio for solutes present in the experimental sample.^21,34-37^ The frictional ratio (*f/f_0_*) is a numerical descriptor of a particle’s anisotropy, such that as the value increases from 1.0, the molecule becomes more asymmetric, moving from spherical to globular, and then to rod-like, with most proteins falling between 1-4.^31,32^ As before,^21^ this modeling method requires the experimental determination of the partial specific volumes 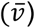 of each of the chromatin fibers analyzed (**Fig. S1**).

The 2DSA/GA-MC analysis indicated that the molecular weights of both WT and H4K16Q arrays were approximately 2.5 MDa, indicating the presence of ~11 nucleosomes on the 12mer templates (**Fig. 2d**). The addition of Sir3 to WT arrays led to an increase in molecular weight to 4.9 Mda, consistent with the binding of 22 molecules of Sir3 (113 kDa each), or a stoichiometry of two Sir3 molecules per nucleosome. In accordance with low levels of binding, the increase in molecular weight of H4K16Q arrays upon Sir3 addition indicated the presence of only ~9 molecules of Sir3. In contrast, the addition of Sir2/4 complex to H4K16Q arrays led to an increase in molecular weight consistent with 22 complexes (of 215 MDa each) bound per array, whereas the increase in molecular weight on WT arrays reflected only 4 complexes bound per array. This difference may indicate Sir2/4 binding is more stable to H4K16Q nucleosomes, which mimic Sir2/4’s natural substrate, H4K16-acetyl, but which cannot be deacetylated by Sir2. This is consistent with a previous study which found Sir2/4 binds with greater affinity to acetylated nucleosomes in the absence of NAD^+^.^27^ When the Sir2/4 complex was added with Sir3, the molecular weight of WT arrays increased to 7.3 MDa, which is consistent with the binding of 22 molecules of Sir3 and 11 molecules of Sir2/4, or a stoichiometry of 2 Sir3 and 1 Sir2/4 per nucleosome. Interestingly, the addition of all three SIR proteins to H4K16Q arrays yielded a similar molecular weight (~7.2 MDa), suggesting that the stoichiometry of SIR proteins is not sensitive to the integrity of H4K16.

Despite a nearly identical increase in molecular weight on addition of SIR proteins, WT arrays sedimented more rapidly than H4K16Q arrays. This suggests that WT arrays adopt a more compacted structure than H4K16Q arrays. This view is reflected in the frictional ratio (*f/f_0_*), as the addition of Sir3 to wildtype arrays increased *f/f_0_* from 2.0 to 2.3, and the further addition of Sir2/4 increased *f/f_0_* to 2.9 (**Fig. 2D**). This suggests a more globular, asymmetric shape of the SIR chromatin fibers compared to nucleosomal arrays or arrays that contain only Sir3. In contrast, the addition of all three SIR proteins to H4K16Q arrays led to an increase of *f/f_0_* from 2.1 to 4.0, indicative of a highly asymmetric, extended conformation. This interpretation is reinforced by the experimentally determined values for the partial specific volumes (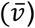) of these fibers, as the wildtype SIR fiber adopted a higher, protein-like value of 0.776 ml/g, whereas the 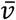 of the SIR H4K16Q fiber remained a low, nucleic acid-like value of 0.679 ml/g, which could reflect the presence of exposed linker DNA in the structure (**Fig. 2d** and **S1**).

### The compaction state of SIR chromatin requires a Sir3-Sir4 interaction

Sir4 contains a C-terminal coiled-coil domain that is required for both Sir4 dimerization and interaction with Sir3. Point mutations in this Sir4 domain have been identified (e.g. Sir4I1311N) that eliminate *in vitro* interactions between Sir4 and Sir3, and *sir4I1131N* disrupts recruitment of Sir3 to silencers *in vivo*.^25,28,29^ To address the contribution of this Sir4-Sir3 interaction on chromatin fiber assembly and dynamics, the Sir2/Sir4 complex was purified from a yeast strain harboring the *sir41131 N* allele, and the interactions of this complex with Sir3 and nucleosomal arrays were first analyzed by EMSA (**Fig. 3a**). On WT chromatin, the Sir2/Sir4113N complex appeared to bind well with Sir3 to form a chromatin fiber with a discrete mobility. However, binding of Sir3 and Sir2/Sir41131N was less effective on the H4K16Q arrays, and the complexes were more diffuse and perhaps less stable.

**Figure 3.**
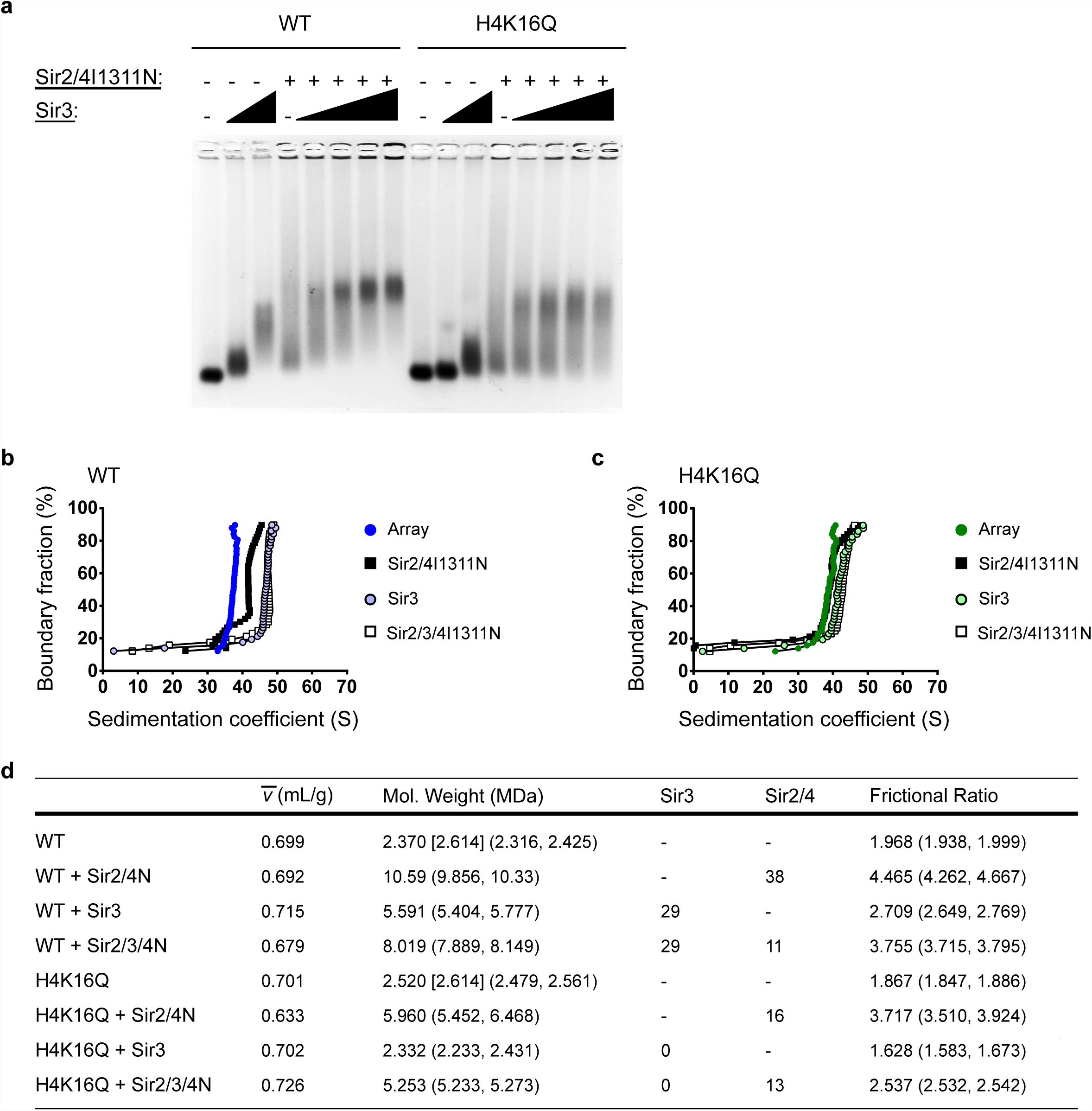
SIR-mediated compaction requires the interaction between Sir3 and Sir4. **(a)** EMSA of Sir3 titrated onto WT and H4K16Q arrays in the absence or present of Sir2/4I1311N. **(b-c)** vHW plots of Sir3, Sir2/4I1311N, and Sir3 and Sir2/4I1311N complex added to WT and H4K16Q arrays. **(d)** 2DSA/GA-MC modeling results of the sedimentation data in **(b-c)**. Numbers in brackets represent expected molecular weights. Numbers in parentheses are 95% confidence intervals. Stoichiometries upon addition of Sir2/3/4 are speculative.

To address the possibility that loss of the Sir3-Sir4 interaction alters the solution dynamics of SIR chromatin fibers, WT and H4K16Q fibers bearing Sir411311N were analyzed by SV-AUC and 2DSA/GA-MC modeling (**Fig. 3b-d**). Two results are apparent from the data. First, the Sir41131N substitution appeared to disrupt the stable binding of Sir3 to the H4K16Q arrays, with no additional increase in molecular weight detected upon Sir3 addition to H4K16Q arrays with Sir2/4 by 2DSA/GA-MC modeling (**Fig. 3d**). Secondly, on WT arrays the Sir3/Sir2/Sir41131N fibers had a molecular weight similar to that of a wildtype SIR chromatin fiber (~8MDa), suggesting a normal subunit stoichiometry. This result was surprising, as the Sir3/Sir2/Sir41131N fibers showed a similar sedimentation profile to fibers assembled with only Sir3 (**Fig. 3b**). These results are explained by the fact that the *f/f_0_* ratio was increased dramatically by the Sir41131N substitution, from 2.0 to 3.8. Similarly, the 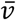 value decreased from 0.715 for Sir3 arrays to 0.679 for Sir3/Sir2/Sir41131N arrays (**Fig. 3d** and **S2**). Interestingly, an *f/f_0_* of 3.8 and 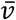 of 0.679 are nearly identical to the values for a SIR H4K16Q fiber, consistent with a more de-condensed structure. These results indicate that the Sir41131N substitution does not disrupt the binding stoichiometry of SIR proteins to WT arrays, but that the interaction between Sir3 and Sir4 is essential for organizing fibers into compact structures.

### SIR proteins compact WT but not H4K16Q arrays

In order to directly observe the structural differences between SIR chromatin fibers, we visualized fibers by atomic force microscopy (AFM). Given the large molecular weight of the SIR complex, we used nucleosomal array substrates reconstituted on DNA templates harboring 36 tandem copies of the 601 nucleosome positioning sequence to visualize the underlying chromatin structure more clearly (**Fig. 4**). As an initial calibration of chromatin folding, WT arrays were initially imaged in low salt buffer in the absence and presence of 1 mM MgCl_2_ which induces folding of extended fibers (-Mg^++^) into condensed 30 nm fibers (+Mg^++^),^38^^-^^40^ with a proportional increase in height from 1.88 to 5.43 nm (**Fig. 4a**). In buffer containing moderate salt (**Fig. 4b**), both WT and H4K16Q arrays adopted a zig-zag structure consistent with an intermediate folded state with a height of approximately 1.9 nm.^21,41^ The addition of Sir3 (**Fig. 4c** and **4b**) increased the height of WT arrays to 4.06 nm, whereas addition of Sir3 to H4K16Q arrays increased the height only slightly to 2.29 nm, corresponding to robust binding to WT but not H4K16Q arrays. Consistent with our previous study,^21^ Sir3 binding appeared to occlude linker DNA (as evidenced by the loss of a beads-on-a-string structure), but Sir3 fibers remained significantly de-condensed compared to the 30 nm fibers shown in **Fig 4a**. In contrast, addition of the full complement of SIR proteins to wildtype arrays led to formation of a more compacted structure, with the height increasing to 5.28 nm (**Fig. 4e**). Notably, this structure is more compact that fibers formed with just the Sir2/Sir4 complex (**Fig. 4d**). Interestingly, addition of all three SIR proteins to H4K16Q arrays increased the height to only 4 nm, and these SIR fibers were clearly more extended and individual nucleosomes could still be identified, suggesting that while SIR proteins bound at similar levels to H4K16Q nucleosomes, they were unable to occlude linker DNA and form a compact structure.

**Figure 4.**
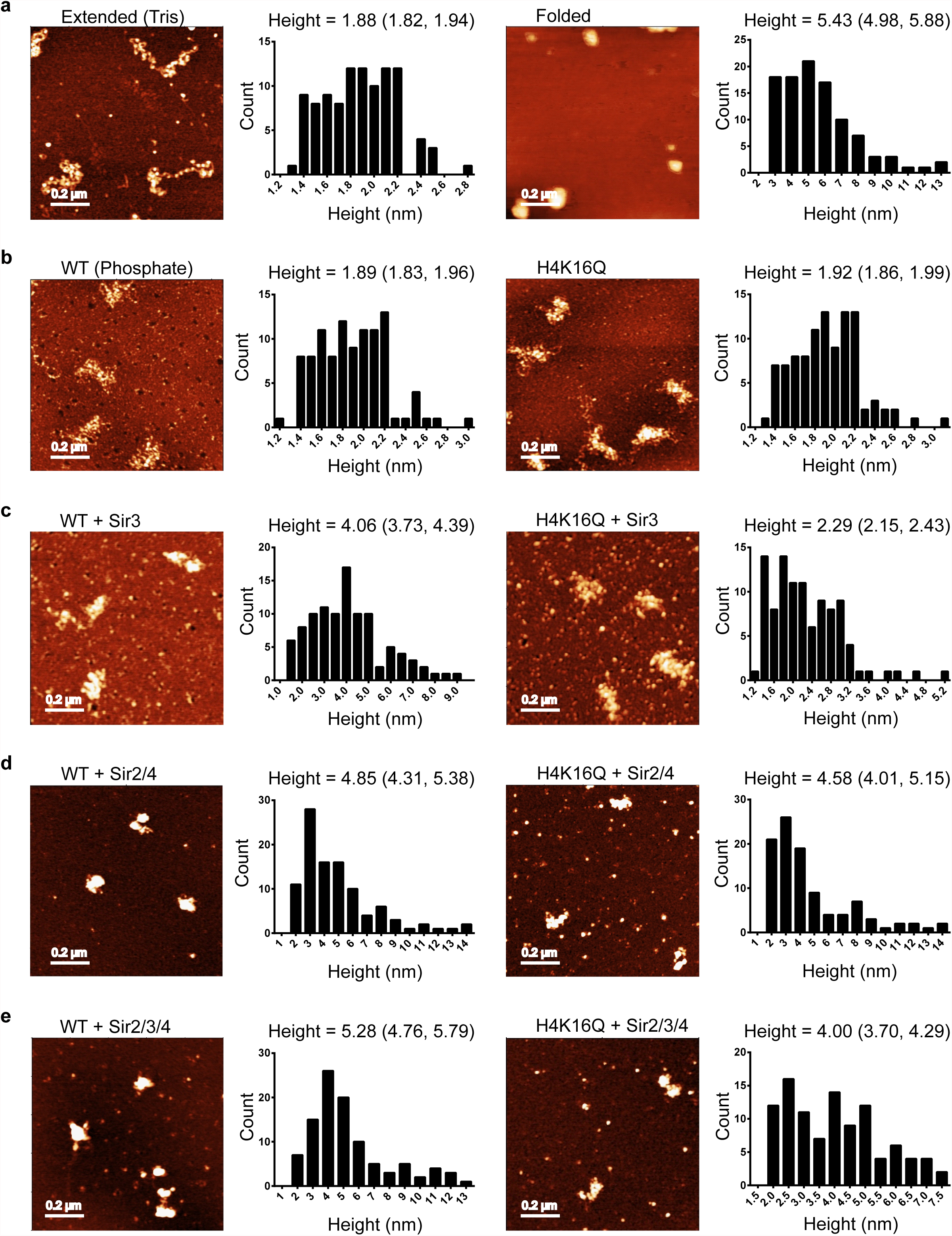
SIR heterochromatin is compact. **(a)** AFM images of WT 601-177-36 arrays in low salt Tris buffer in the absence (left) or presence (right) of 1 mM MgCl_2_. Histograms are of 100 individual measurements. Mean height and 95% confidence intervals are shown above. **(b-e)** WT and H4K16Q 601-177-36 arrays in sodium phosphate buffer incubated with indicated SIR proteins.

## Discussion

Heterochromatin fibers are believed to be highly compact structures that block DNA accessibility to DNA binding transcription factors and components of the recombinational repair machinery.^2,3,17^ The *in vitro* assembly of budding yeast SIR chromatin fibers yields these expected functional properties – such fibers hinder access of restriction enzymes, repress *in vitro* transcription by RNA polymerase II, and block early steps of *in vitro* recombinational repair.^20,25,26^ What has been limiting, however, is evidence that SIR chromatin fibers form condensed structures consistent with these repressive properties. Previous studies have yielded conflicting results whereby addition of SIR proteins to nucleosomes either led to formation of long, extended filaments,^19,42^ or to fibers where SIR proteins appeared to aggregate nucleosomes.^25^ In both previous cases, SIR chromatin fibers were not analyzed in solution but visualized after immobilization on EM grids. Here, we have undertaken the first solution analysis of the conformational dynamics of SIR chromatin fibers. Using a combination of SVAUC and molecular modeling, our results demonstrate formation of nearly homogeneous SIR chromatin fibers that are compact, not extended. Furthermore, this compact structure requires the integrity of H4K16, as well as a physical interaction between Sir3 and Sir4. Finally, the compact structure indicated by the solution methods is confirmed by AFM analysis.

### Multiple roles for Sir3-Sir4 interactions

Previous studies have shown that the Sir4I1311N substitution eliminates interactions between Sir3 and Sir4 *in vitro* and disrupts the recruitment of Sir3 to silencing domains *in vivo*.^25,28,29^ Furthermore, we found that Sir4I1311N disrupts the binding of Sir3 to H4K16Q nucleosomal arrays that harbor Sir2/Sir4, consistent with a previous study showing a similar impact on Sir3 binding to arrays containing H4K16A.^25^ These results are all consistent with a recruitment role for the Sir3-Sir4 interaction surface,^9,29^ and suggest that the presence of Sir3 on Sir2/4-bound chromatin lacking H4K16 is due to Sir3 binding to Sir4 and not to the H4 tail. Surprisingly, we found that the Sir4I1311N substitution eliminated the formation of compact SIR chromatin fibers, even though it had no apparent impact on the stoichiometry of SIR proteins on WT arrays. This suggests that interactions between Sir4 and Sir3 are key for the proper organization of SIR proteins on nucleosomal arrays, ensuring that Sir subunits are oriented in a manner that leads to a functional, condensed structure. As the Sir4 coiled-coil domain binds neither DNA nor nucleosomes,^29^ it likely serves as a bridge, perhaps by mediating cross-nucleosomal Sir3-Sir3 interactions as well as essential Sir3-Sir4 interactions that drive fiber condensation.

### The shared role of Sir3 and Sir2/4 on heterochromatin formation and function

*In vivo*, overexpression of Sir3 can create extended, transcriptionally silent chromatin domains that are severely depleted of Sir2 and Sir4.^10,43^ Likewise, Sir3 addition to nucleosomal arrays is sufficient to inhibit early steps of recombinational repair, though this inhibition is strengthened by further addition of Sir2 and Sir4.^20^ These data are consistent with the view that Sir3 is sufficient to create a partial, perhaps weaker form of heterochromatin fiber. Notably, Sir3 chromatin fibers are not highly condensed,^21^ but Sir3 does appear to stabilize nucleosomes,^26^ perhaps by mediating contacts between arginines 17 and 19 of the H4 tail and DNA.^24^ Additionally, the occusion of DNA linkers in Sir3 chromatin fibers requires its dimerization domain,^21^ consistent with Sir3 dimerization bridging adjacent nucleosomes.^44^ We propose that the Sir2/Sir4 complex reinforces the silencing properties of SIR heterochromatin by folding the SIR chromatin fiber into an inaccessible, condensed structure. Our analysis of SIR protein stoichiometry suggests a model in which one molecule of Sir2/4 binds primarily within each nucleosome linker and two molecules of Sir3 bind to each surface of the nucleosome, and fiber compaction is completed via cross-nucleosomal Sir3-Sir4 and Sir3-Sir3 interactions (**Fig. 5**). The resulting SIR chromatin fiber would be characterized by both stabilized nucleosomes and occlusion of DNA linkers, leading to compact structure that effectively represses DNA-templated reactions.

**Figure 5.**
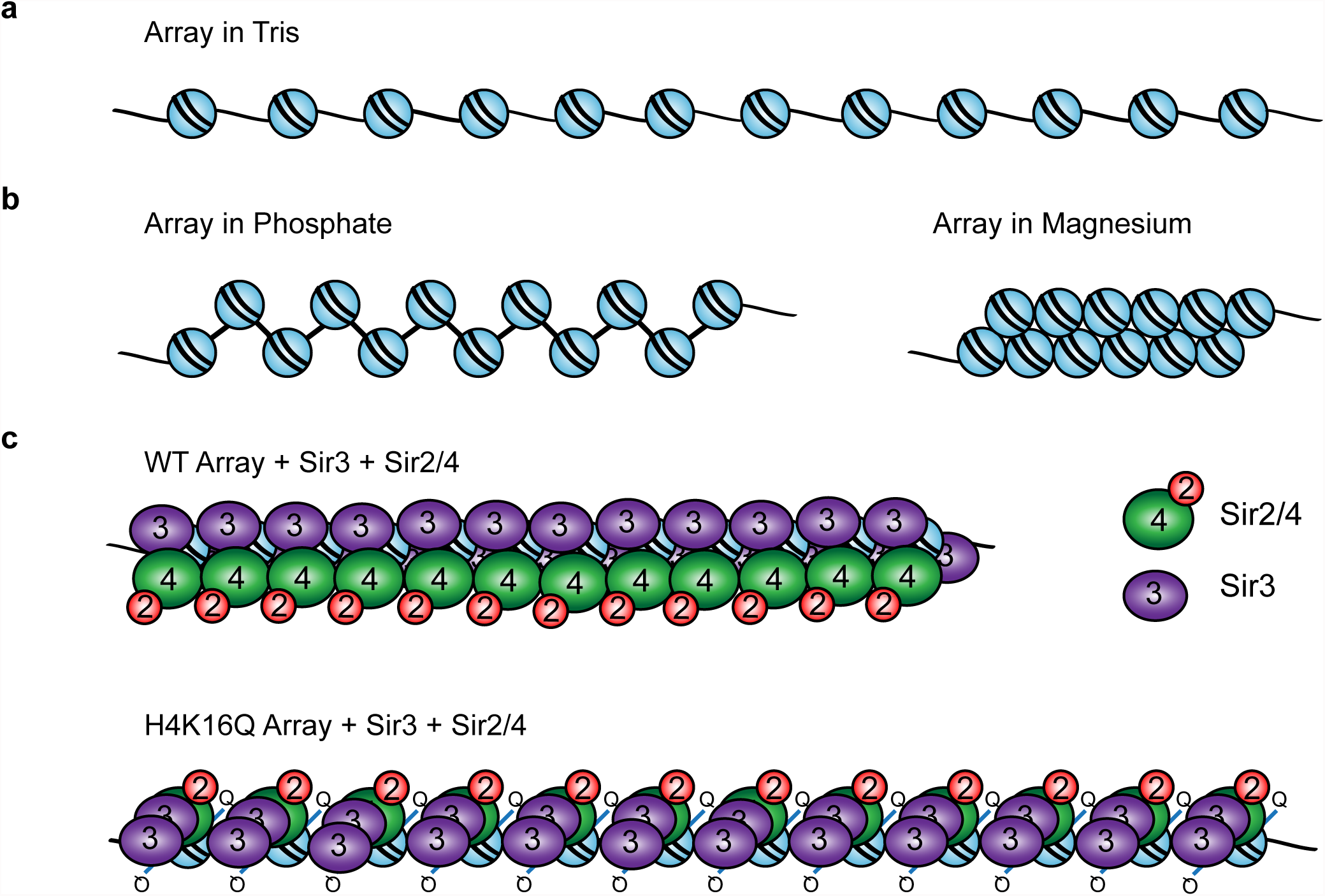
Model for a SIR chromatin fiber. **(a)** Diagram of a 12-mer array in low-salt Tris buffer. **(b)** Arrays in 20 mM phosphate buffer pH 8.0 (containing ~40 mM Na^+^) are partially folded. Arrays in 1 mM MgCl_2_ buffer fold into 30 nm fibers. **(c)** SIR proteins bind and condense WT arrays, though to a lesser extent than 30 nm fibers, with two molecules of Sir3 and likely one molecule of Sir2/4 per nucleosome. Although Sir proteins also bind H4K16Q arrays, linker DNA is not occluded.

## Acknowledgements

We thank members of the Peterson lab for helpful discussions. This work was supported by grants from the NIH to C.L.P (GM54096) and S.A.W. (R01AI117839), and a grant from the NCI to S.L. (U54 CA143862).

## Author Contributions

AFM experiments were performed by S.S., and all other experiments and reagent prepration were performed by S.G.S. M.F.B. and S.A.W. created the 601-177-36 plasmid. Data analysis was performed by S.G.S., S.S., S.L., and C.L.P. The manuscript was prepared by S.G.S. and C.L.P.

## Competing Financial Interests

None

## Materials and Methods

### Proteins

FLAG-tagged Sir3 protein and TAP-tagged Sir2/4 complex were individually overexpressed and affinity purified from yeast. Briefly, yeast cultures transformed with plasmids contained tagged proteins under a galactose-inducible promoter were grown to OD 0.6 and induced with 2% galactose for 5 hours. Cultures were pelleted, resuspended in E Buffer (20 mM HEPES pH 7.4, 350 mM NaCl, 10% glycerol, 0.1% Tween 20, and protease inhibitors), and frozen in liquid nitrogen. Pellets were ground using a cold mortar and pestle with frequent additions of liquid nitrogen until approximately 50% of cells appeared lysed under a microscope. Cells were incubated on ice in E buffer for 30 min, then spun at 3,000 rpm for 15 minutes to remove debris. Supernatant was spun down at 40,000 rpm for 1 hour, then the aqueous layer was removed from the lipid layer using a syringe. For Sir3 purification, lysate was incubated with anti-Flag resin from Sigma for three hours at 4°C. Resin was washed in E buffer, then Sir3 was eluted in batch via four 30 minute incubations of resin with E Buffer containing 100 µg/mL 3xFLAG peptide from Sigma. For Sir2/4 purification, lysate was incubated with IgG resin for 2 hours, washed in E buffer, then eluted in batch via the addition of purified TEV protease overnight. Eluted Sir2/4 was then bound in batch to Calmodulin resin for 2 hours in the presence of Ca^2+^, washed in E buffer, and eluted with EGTA. Concentrations were determined by comparison to known concentrations of BSA electrophoreses on the same Coomassie-stained SDS-PAGE gel. The Sir2/4I1311N plasmid was generated by site-directed mutagenesis, and purified as above. All Sir2/4 was dialyzed into 20 mM sodium phosphate buffer pH 8.0 prior to use in order to maintain moderate concentrations of salt across experiments. Recombinant *Xenopus laevis* histones were expressed in BL21 cells, purified, and assembled into histone octamers according to standard protocols.

### DNA

The 601-177-12 nucleosomal array template containing twelve copies of the Widom 601 nucleosome positioning sequence was digested from its plasmid backbone using EcoRV and purified by size-exclusion chromatography. The 601-177-36 fragment was generated by HaeII and XbaI digestion of the 601-177-36 plasmid and purified as above. The 36×601 multinucleosomal array sequence was generated in three assembly steps. First, 12×601 sequences were generated via golden gate assembly^45,46^ from individual monomers. Briefly, the 601 nucleosome positioning sequence was first PCR amplified with primers carrying a unique barcode sequence and BsaI cleavage site. Twelve of these 601 PCR amplicons were assembled via golden gate as an array (12×601) into the pFUS-A backbone^47^ via BsaI digestion and ligation to generate three different arrays (MA-1, MA-2 & MA-3). Next, two 12×601 sequences were concataned to generate 24×601 sequences as follows: one of the 12×601 plasmids (MA-1) was digested with SpeI and SphI and the backbone was recovered; a second 12×601 plasmid (MA-2) was digested with XbaI and SphI and the released 12×601 array was recovered; then, these two components were ligated. This process was repeated using the MA-3 12×601 array to generate the 36×601 sequence.

### Nucleosomal array assembly

Nucleosomal arrays were assembled by combining recombinant histone octamers and 601-177-12 or 601-177-12 DNA template at varying molar ratios of octamer to nucleosome positioning sequence in 2 M NaCl, and step-wise salt dialysis was performed until completion into 20 mM sodium phosphate pH. 8.0 with 0.1 mM EDTA. Array saturation was determined by ScaI digestion followed by analysis via native PAGE and by SV-AUC.

### EMSA

300 ng WT or H4-K16Q nucleosomal array was combined with Sir2/4 at a ratio of 1, 2, or 3 molecules per nucleosome to a final concentration of 10 ng/ul array and 5% glycerol. For combined Sir2/4 and Sir3 EMSA, Sir2/4 or Sir2/4I1311N was added at 2 molecules per nucleosome, and Sir3 was titrated at 1, 2, 4, and 6 molecules per nucleosome. Binding reactions were incubated at room temperature for 30 minutes, run on 1% TBE agarose gels, and stained with ethidium bromide.

### SV-AUC

SV-AUC was carried out using 400 µl sample loaded into two-sector Epon centerpieces in an An60 Ti rotor in a Beckman Optima XL-I analytical ultracentrifuge, and run at 20°C. Measurement was completed in intensity mode. Nucleosomal arrays were run at 10 ng/ul concentrations with the indicated amounts of Sir3 or Sir2/4 at 20,000 RPM, and were measured at 215 nm (for arrays alone) or 260 nm (for samples containing SIR proteins). For experiments containing all three SIR proteins, Sir2/4 was added first, followed by array, followed by Sir3. For 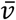 determination, three preparations of sample were run as above, with 0, 25, or 50% H_2_^18^O (obtained from Cambridge Istotope Laboratories, Andover, MA) added in place of H_2_^16^O. The obtained S values were then plotted as a function of solvent densities, linear regression was performed, and the 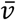 was calculated by dividing the slope of the resulting line by the y-intercept. Solvent densities and viscosities were obtained from the literature. Linear regression was performed using GraphPad Prism software.

### 2DSA/GA-MC

All SV-AUC data were analyzed using UltraScan3 software, version 3.3 and release 1977 (http://www.ultrascan3.uthscsa.edu/index.php), and fitting procedures were completed on XSEDE clusters at the Texas Advanced Computing Center (Lonestar, Stampede) and at the San Diego Supercomputing Center (Trestles) through the UltraScan Science Gateway (https://www.xsede.org/web/guest/gateways-listing). Raw intensity data were converted to pseudo-absorbance by using the intensity of the air above the meniscus as a reference and edited. As previously described,^21^ partial specific volumes (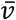) of each of the chromatin fibers were determined experimentally (**Fig. S1**). Next, 2DSA was performed to subtract time-invariant noise and the meniscus was fit using 10 points in a 0.05 cm range. Arrays were fit using an S range of 5-60 S, an *f/f_0_* range of 1-10 with 100 grid points for each, 10 uniform grid repetitions, 400 simulation points, and meniscus fitting within a 0.6 cm range with 10 points. 2DSA was then repeated at the determined meniscus to fit radially-invariant and time-invariant noise together using 5 iterations. vHW analysis was completed using these noise subtraction profiles to determine S. Where indicated, GA was initialized by binning major solutes in the 2DSA dataset, and run via LIMS. Major solutes from GA analysis were then binned and run again using GA with 50 MC iterations.

### AFM

For atomic force microscopic experiments, an Agilent AFM 5500 instrument and silicon nitride cantilevers were used (force constant 25-75 N/m, resonant frequency 332 kHz). Imaging was done in air using the acoustic AC mode with an amplitude of ~10 nm and a set-point reduction of about 10%, scanning at 1 line per second. Immobilization of chromatin arrays on mica surface was done as follows. First, Sir3 or Sir2/4 was added to phosphate buffer followed by addition of 10 ng/ul chromatin array and mixed gently, maintaining a ratio of 4 Sir3 or Sir2/4 molecules/nucleosome. For imaging with both Sir3 and Sir2/4, Sir2/4 was added first, then arrays, followed by Sir3, at a ratio of 2 Sir3’s and 2 Sir2/4’s per nucleosome. After 30 minutes, 0.5% glutaraldehyde solution (1 µL) was added to this mixture for crosslinking and incubated for 10 minutes. APTES was deposited on freshly cleaved mica substrate using vapor deposition. The crosslinked chromatin solution was diluted to 1 ng/µL and 3 µL was added to this APTES modified mica surface and after 5 minutes the surface was cleaned three times using 400 µL of buffer solution, dried carefully using argon gas and immediately used for imaging. To image only chromatin arrays, the first mixing step with SIR proteins was omitted, and imaging was carried out in the indicated buffer. Nucleosomal heights were measured using Gwyddion software, and then mean heights were computed for each experimental condition.

## Figure Legends

**Supplementary Figure 1.**
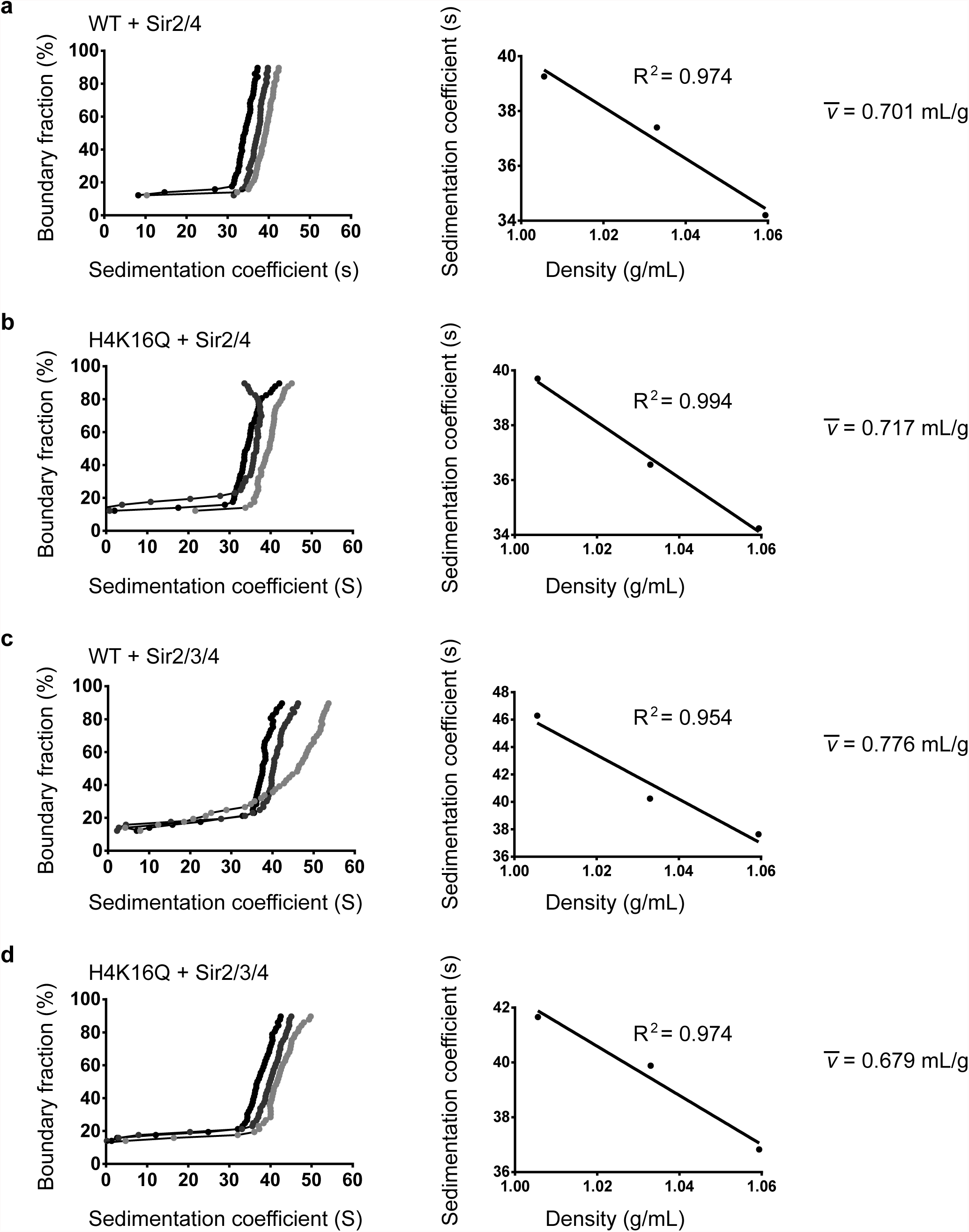
The partial specific volume of WT and H4K16Q arrays with Sir2/4. **(a-d)** vHW plots showing the sedimentation of molecules in 0% (light gray), 25% (dark gray), and 50% H2O18 (black) and plots of sedimentation coefficient vs. density. The 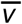 is calculated by dividing the slope of the fit line by the y-intercept.

**Supplementary Figure 2.**
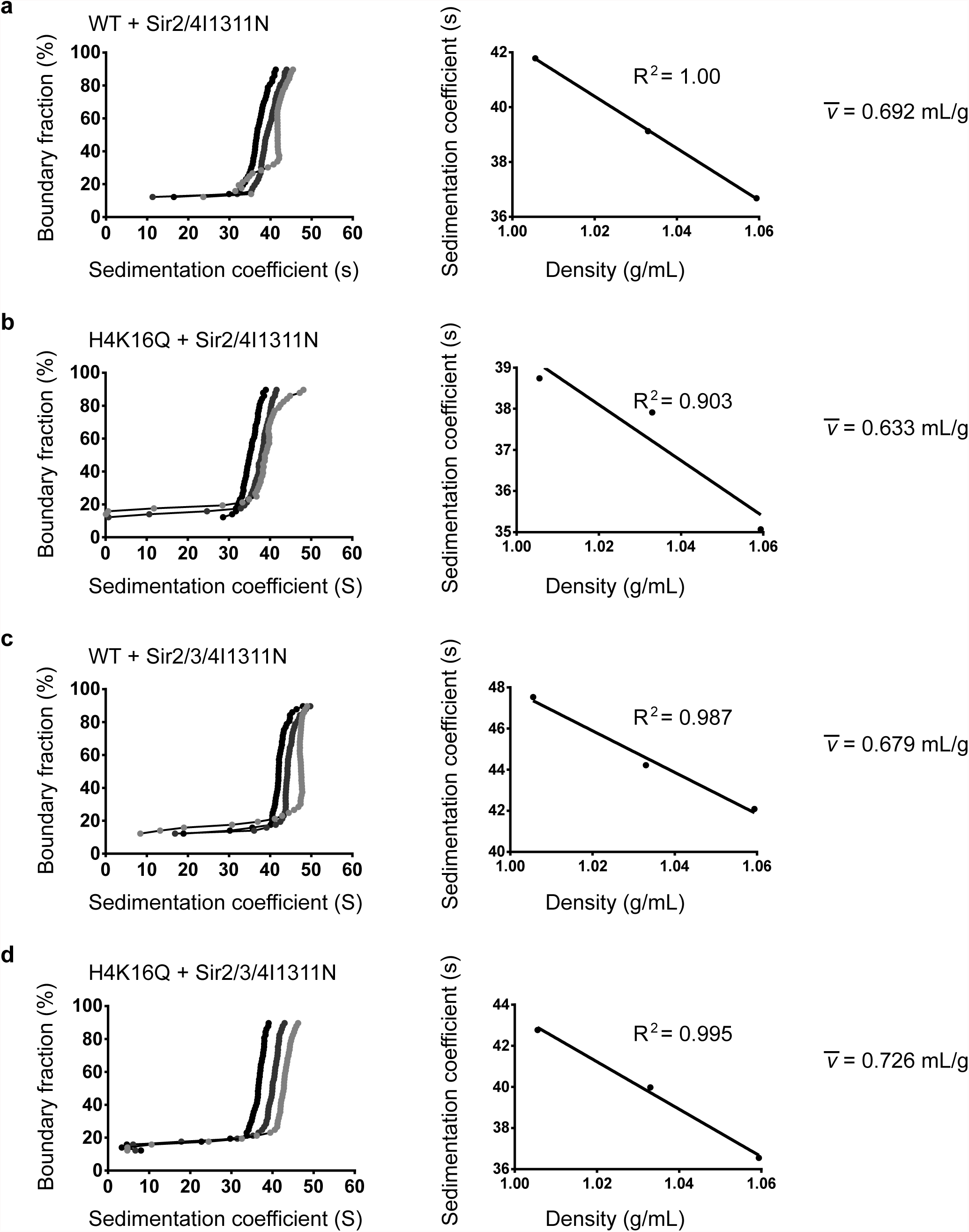
The partial specific volume of WT and H4K16Q arrays with Sir2/4I1311N. **(a-d)** vHW plots showing the sedimentation of molecules in 0% (light gray), 25% (dark gray), and 50% H2O18 (black) and plots of sedimentation coefficient vs. density. The 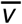 is calculated by dividing the slope of the fit line by the y-intercept.

